# Maternal Dppa2 and Dppa4 are dispensable for zygotic genome activation but important for offspring survival

**DOI:** 10.1101/2021.09.13.460183

**Authors:** Oana Kubinyecz, Fatima Santos, Deborah Drage, Wolf Reik, Melanie A Eckersley-Maslin

## Abstract

Zygotic Genome Activation (ZGA) represents the initiation of transcription following fertilisation. Despite its importance in shifting developmental control from primarily maternal stores in the oocyte to the embryo proper, we know little of the molecular events that initiate ZGA *in vivo*. Recent *in vitro* studies in mouse embryonic stem cells (ESCs) have revealed Developmental Pluripotency Associated 2 and 4 (Dppa2/4) as key regulators of ZGA-associated transcription. However, their roles in initiating ZGA *in vivo* remain unexplored. We reveal Dppa2/4 proteins are present in the nucleus at all stages of preimplantation development and associate with mitotic chromatin. We generated single and double maternal knockout mouse models to deplete maternal stores of Dppa2/4. Importantly, while fertile, Dppa2/4 maternal knockout mice had reduced litter sizes, indicating decreased offspring survival. Immunofluorescence and transcriptome analyses of 2-cell embryos revealed while ZGA took place there were subtle defects in embryos lacking maternal Dppa2/4. Strikingly, heterozygous offspring that inherited the null allele maternally had higher preweaning lethality than those that inherited the null allele paternally. Together our results show that while Dppa2/4 are dispensable for ZGA transcription, maternal stores have an important role in offspring survival, potentially via epigenetic priming of developmental genes.

## Introduction

One of the first developmental milestones following fertilisation is transcriptional activation of the embryonic genome. This process, termed zygotic genome activation (ZGA) is part of the maternal to zygotic transition and considered essential for development to progress. It occurs in two phases: a minor wave at the late zygote/early 2 cell stage and a major wave at the middle-to-late 2-cell stage in mouse (4- to 8-cell stage in humans). At the time of ZGA, the preimplantation embryo undergoes dramatic epigenetic reprogramming (reviewed in (Eckersley-Maslin et al., 2018)), however we have limited understanding of the maternal factors that regulate ZGA and epigenetic reprogramming *in vivo* mostly due to complex genetics and limited cell numbers.

More recently, the field has modelled ZGA using a spontaneously occurring rare subpopulation of mouse embryonic stem cells (ESCs), termed 2C-like cells (Macfarlan et al., 2012). These cells share many of the epigenetic and transcriptional features of the 2-cell embryo (reviewed in (Eckersley-Maslin et al., 2018; Genet and Torres-Padilla, 2020)), and recent studies have exploited these similarities to screen for regulators of ZGA-associated transcription *in vitro* (Alda-Catalinas et al., 2020; Eckersley-Maslin et al., 2019; Grow et al., 2021; Hu et al., 2020; Rodriguez-Terrones et al., 2018; Zhao et al., 2018). From these and other studies, the dimerising nuclear proteins *Developmental Pluripotency Associated 2* (Dppa2) and *4* (Dppa4) were identified as potent inducers of the 2C-like state (De Iaco et al., 2019; Eckersley-Maslin et al., 2019; Yan et al., 2019). In ESCs, Dppa2/4 directly bind to and induce transcription of the *Dux* transcription factor gene which in turn regulates expression of downstream ZGA targets (De Iaco et al., 2017; Hendrickson et al., 2017; Whiddon et al., 2017). Additionally, Dppa2/4 also regulate other transcripts outside of the Dux network, including LINE-1 elements (De Iaco et al., 2019; Gretarsson and Hackett, 2020) and a subset of bivalently marked developmental genes (Eckersley-Maslin et al., 2020; Gretarsson and Hackett, 2020).

Dppa2/4 are present in mature oocytes, further upregulated during ZGA and persist throughout preimplantation development only to be silenced by DNA methylation upon gastrulation at E6.5 (Eckersley-Maslin et al., 2019; Maldonado-Saldivia et al., 2007). While single and double Dppa2/4 zygotic knockout mice survive embryogenesis, they succumb to lung and skeletal defects shortly after birth despite not being expressed in these tissues (Madan et al., 2009; Nakamura et al., 2011). This suggests they act by priming the epigenetic landscape earlier in development to enable successful development to take place (reviewed in (Eckersley-Maslin, 2020; Watabe, 2012; Zlotorynski, 2020)). However, the role Dppa2/4 have in preimplantation embryos remains elusive as current zygotic knockout models are confounded by maternal deposits of these proteins, and homozygous Dppa2/4 knockout mice have near 100% lethality.

To assess the importance of maternal Dppa2/4 in early embryonic development, including ZGA, we generated conditional single and double knockout mouse models to deplete maternal stores of Dppa2/4 in growing oocytes. Maternal knockout Dppa2/4 mice are fertile and give rise to viable offspring. Furthermore, molecular analyses reveal that ZGA takes place in 2-cell embryos derived from maternal knockouts, indicating Dppa2/4 is not required to initiate ZGA *in vivo*. However, heterozygote offspring from maternal knockouts have increased preweaning mortality compared to those from paternal knockouts. Therefore, while dispensable for ZGA to take place, maternal stores of Dppa2/4 have an important role in offspring survival.

## Results

### Dppa2/4 proteins localise to euchromatin in preimplantation embryos and associate with mitotic chromatin

Dppa2/4 transcripts are present in growing and mature oocytes and throughout preimplantation development (Eckersley-Maslin et al., 2019), however protein expression patterns and localisation have not been systematically assessed at these developmental stages. Immunofluorescence staining on wild-type C57Bl/6 embryos derived from natural matings confirmed the presence of Dppa4, but not Dppa2, in MII oocytes and zygotes (Figure 1A). Dppa4 protein was present in both pronuclei by PN3 prior to the minor wave of ZGA (Figure S1A). There was a marked increase in protein levels from the 2-cell stage onwards following DNA replication (Figure S1B) consistent with their transcriptional upregulation during the major wave of ZGA. Nuclear localisation levels of Dppa2/4 remained constant and in all blastomeres from the 2-cell stage through to the morula, and at the blastocyst stage both proteins showed increased signal in the inner cell mass (ICM) compared to the trophectoderm cells. At all stages, there was colocalization of Dppa2 and Dppa4 proteins with each other consistent with them forming heterodimers (Figure 1A).

**Figure 1:**
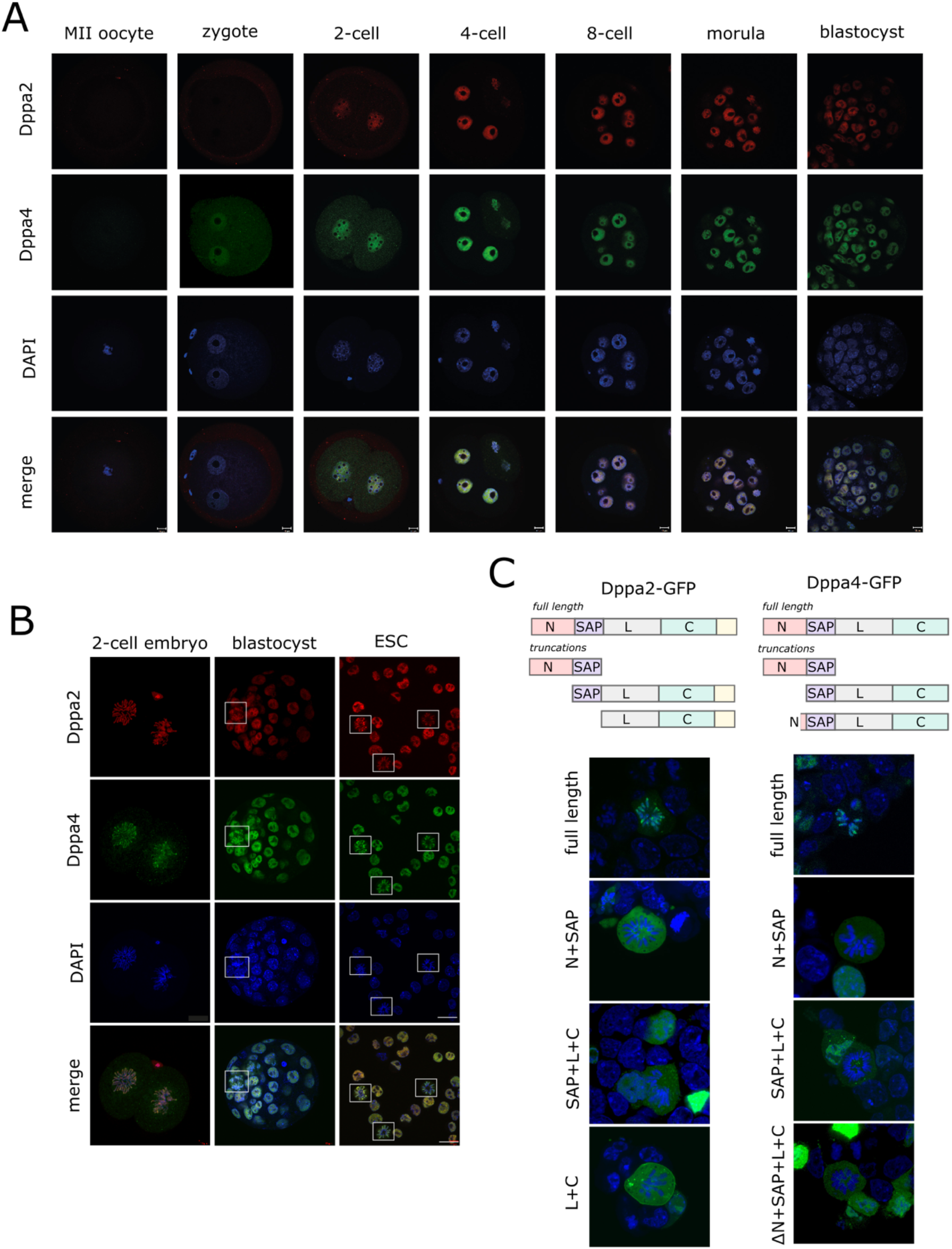
Dppa2/4 localise to euchromatin in preimplantation embryos and associate with mitotic chromatin. (A) Immunofluorescence staining of wild type mouse MII oocytes and preimplantation embryos with Dppa2 (red), Dppa4 (green) and DAPI nuclear stain (blue). Scale bar represents 10µm. (B) Immunofluorescence staining of wild type 2-cell embryos (left), blastocysts (middle) or mouse embryonic stem cells (mESC, right) for Dppa2 (top row, red), Dppa4 (second row, green), DAPI nuclear stain (third row, blue). Mitotic cells are denoted by boxes. (C) Localisaton of Dppa2-GFP (left) or Dppa4-GFP (right) constructs in mouse embryonic stem cells. Full length (top) or truncations (bottom three) were transiently transfected in to cells. Schematic of truncations analysed is shown (not to scale), N-terminus (N), SAP, linker (L) or C-terminus (C) domains are denoted.

Remarkably, we observed strong Dppa2/4 binding to mitotic chromatin in both embryos and embryonic stem cells (ESCs) (Figure 1B), consistent with recent proteomic profiling of mitotic chromatin in ESCs (Djeghloul et al., 2019). To determine if there was a particular region of Dppa2/4 that was responsible for this mitotic binding, we analysed a series of GFP-tagged truncated constructs in ESCs (Figure 1C). Dppa2/4 contain a SAP domain and a C-terminal domain that bind RNA/DNA and histones respectively (Masaki et al., 2010). While full length Dppa2 and Dppa4 bound mitotic chromatin, all truncations analysed lost this ability despite retaining their nuclear localisation in non-mitotic cells. This indicates that multiple regions of the protein, including the SAP domain, N-terminus and C-terminus, are required together for mitotic binding, and that neither domain on its own is sufficient.

### Maternal Dppa2/4 single and double knockout mice are fertile

Having validated the presence of Dppa2/4 protein in preimplantation embryos, we next generated conditional knockout mouse models for *Dppa2/4*. For the single conditional knockouts, LoxP sites were introduced either side of exon 2 of *Dppa2* or *Dppa4* (Figure 2A, supplemental materials and methods). *Dppa2/4* genes are located in tandem on mouse chromosome 16 with no annotated intervening genes. This allowed us to generate a conditional double knockout mouse model by introducing LoxP sites upstream of *Dppa4* exon 2 and downstream of *Dppa2* exon 7 (Figure 2A). Conditional knockout mice (Figure 2B) were crossed with Zp3-Cre males to delete *Dppa2* and/or *Dppa4* in growing oocytes and generate maternal knockouts (denoted here on as Dppa2^m-^, Dppa4^m-^ and Dppa2/4^m-^ for the single and double maternal knockouts respectively).

**Figure 2:**
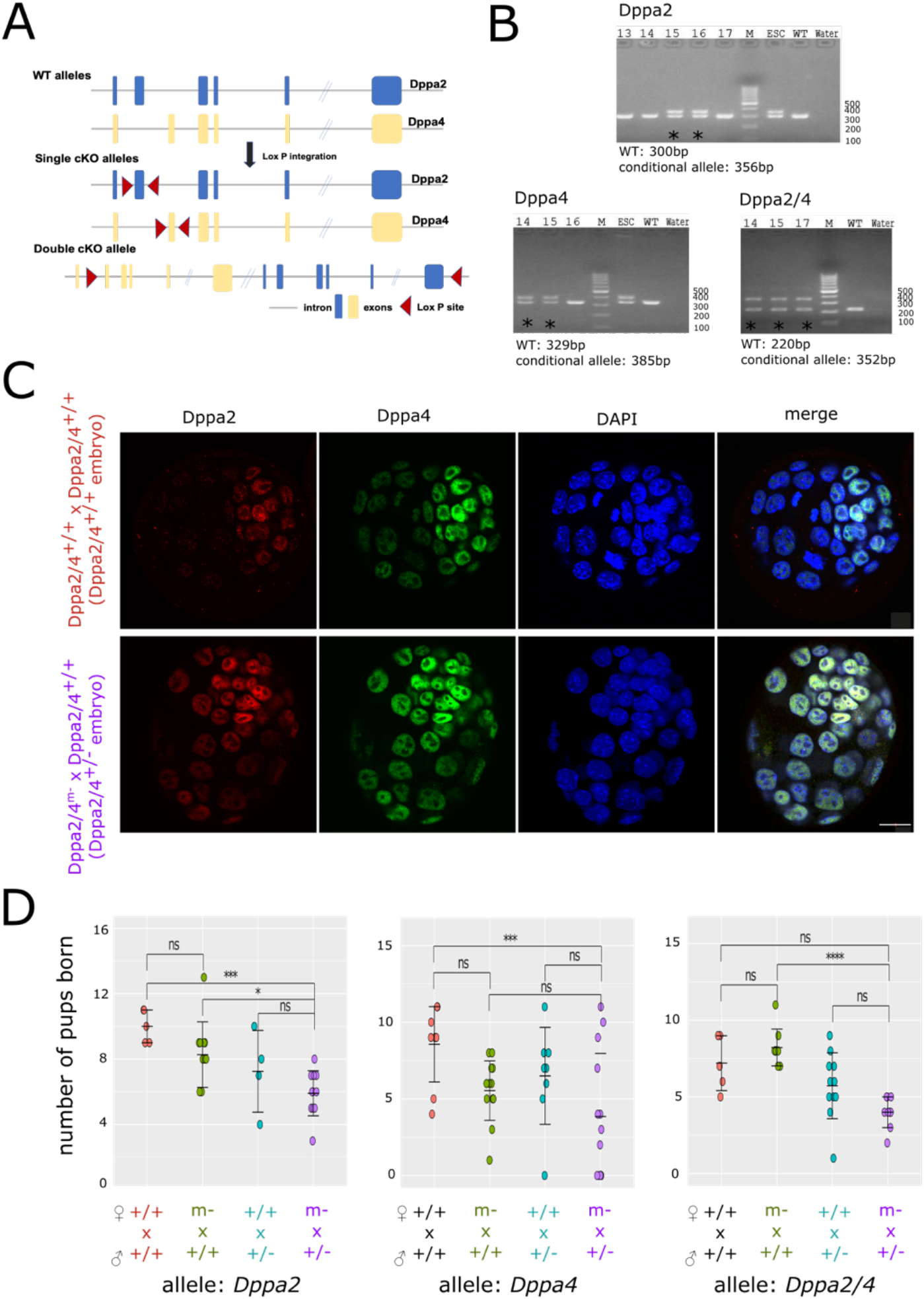
Maternal Dppa2/4 single and double knockout mice are fertile. (A) Schematic of conditional knockout alleles (cKO, not to scale) showing exons (boxes) bounded by LoxP sites (red triangles), Dppa2 allele is shown in blue, Dppa4 allele in yellow. genotyping PCR confirming successful generation of heterozygous mice for Dppa2 (top), Dppa4 (bottom left) or Dppa2/4 (bottom right) conditional allele. * denotes heterozygous founding animals (C) Immunofluorescence staining of blastocysts collected from Dppa2/4^+/+^ (top) or Dppa2/4^m-^ (bottom) females crossed with Dppa2/4^+/+^ fathers for Dppa2 (left, red), Dppa4 (second column, green) and DAPI nuclear stain (third column, blue). Scale bar represents 20µm. (D) litter size for Dppa2 (left), Dppa4 (middle) or Dppa2/4 (right) conditional knockout alleles. Each dot represents an individual litter. Error bars represent mean +/- standard deviation. Statistically significant differences are denoted **p<0.01, ***p<0.001, ****p<0.0001, ns non significant (One-Way Anova with Tuckey multiple comparisons test).

We first assessed whether maternal stores of Dppa2/4 are required to form blastocysts. We crossed control or Dppa2/4^m-^ females with Dppa2/4^+/+^ males and collected the heterozygous Dppa2/4^+/-^ embryos at E3.5. We readily collected phenotypically normal blastocysts from Dppa2/4^m-^ females (Figure 2C). These embryos expressed Dppa2/4 protein from the paternally inherited wild type allele, suggesting ZGA had taken place. Therefore, maternal Dppa2/4 is not required for blastocyst development.

To determine whether maternal Dppa2/4 are required for complete embryonic development, we crossed the Dppa2^m-^, Dppa4^m-^ and Dppa2/4^m-^ with wild type males and monitored litter size and survival. All three genotypes repeatedly gave rise to live litters when crossed with wild type males (Figure 2D). In the case of Dppa2^m-^ and Dppa4^m-^, the resulting litter sizes were smaller, indicating decreased offspring survival. This was exaggerated when crossing single and double maternal knockout females with heterozygote males where there was complete litter loss in some instances (Figure 2D). These results indicate that, while dispensable for blastocyst formation, maternal stores of Dppa2/4 are required for offspring survival and cannot be fully compensated by a WT paternal allele.

### 2-cell embryos from maternal knockout females undergo successful ZGA with subtle differences

While Dppa2/4^m-^ females give rise to offspring, their litters are smaller than their wild type counterparts. Therefore, we analysed 2-cell embryos at a molecular level to determine if there were any ZGA defects. First, we collected 2-cell embryos from WT or maternal knockout females of all three genotypes crossed with WT males and performed immunofluorescence for the MERVL endogenous retrovirus gag protein (Figure 3A) which is used as a marker for ZGA (Kigami et al., 2003; Peaston et al., 2004). As expected, embryos from Dppa2/4^+/+^ x Dppa2/4^+/+^ crosses had high levels of cytoplasmic MERVL gag protein indicating that ZGA had taken place. Heterozygous Dppa2^+/-^, Dppa4^+/-^ and Dppa2/4^+/-^ embryos from maternal knockout females all had reduced yet detectable levels of Dppa2 and Dppa4 proteins (Figure 3A-B), indicating expression from the paternal allele.

**Figure 3:**
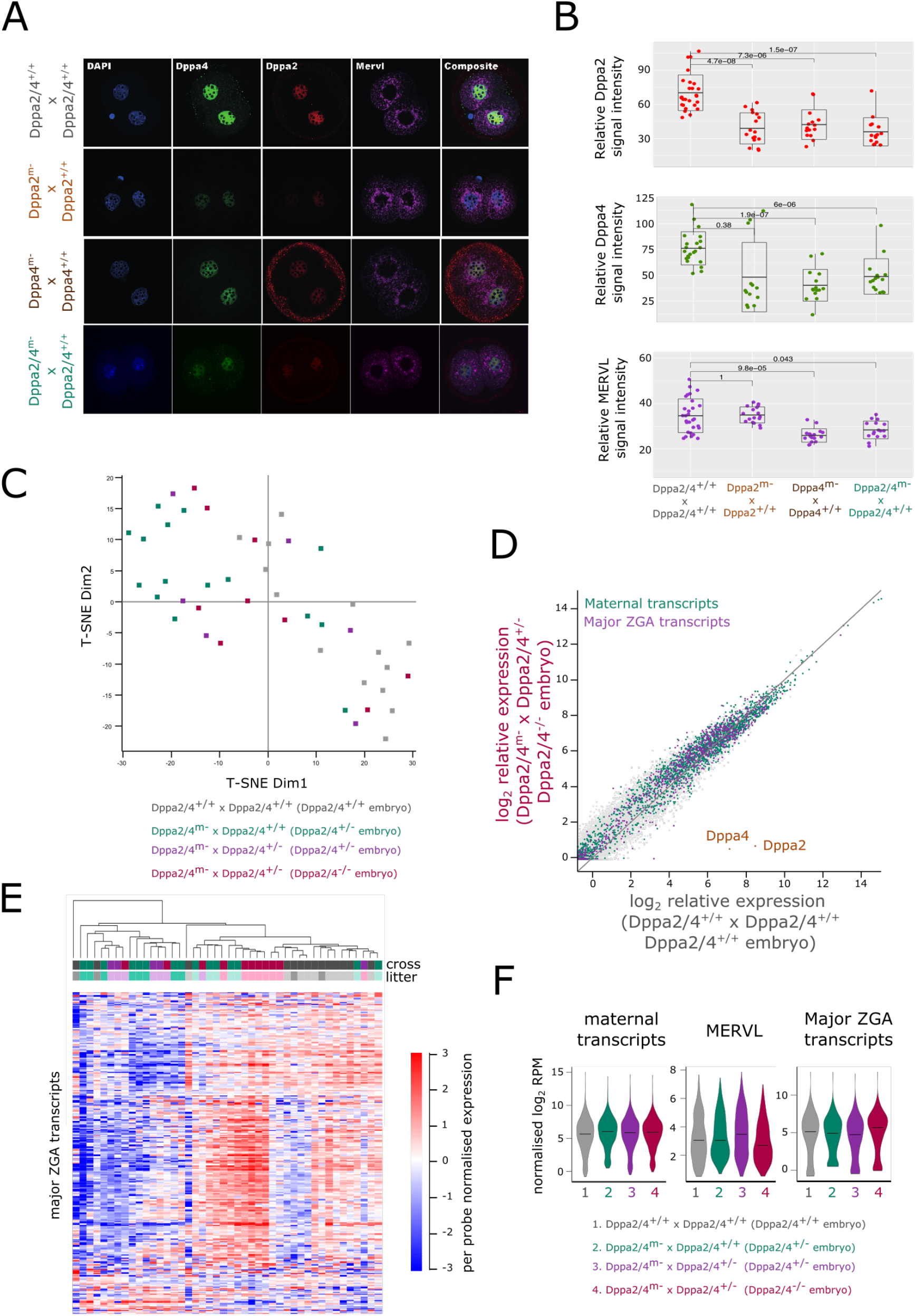
2-cell embryos from maternal knockout females undergo successful ZGA with subtle differences. (A) Immunofluorescence staining of 2-cell embryos derived from Dppa2/4^+/+^, Dppa2^m-^, Dppa4^m-^ or Dppa2/4^m-^ females crossed with wild type fathers, for Dppa2 (red), Dppa4 (green), MERVL (magenta) and DAPI nuclear stain (blue). Scale bar represents 10µm. (B) Relative quantification of MERVL cytoplasmic signal intensity of 2-cell embryos in A. Differences are statistically significant and p-values denoted (Wilcoxon test with Bonferroni correction for multiple comparisons). (C) T-SNE plot of individual 2-cell embryos that are Dppa2/4^+/+^ (grey), Dppa2/4^+/-^ (green, purple) or Dppa2/4^-/-^ (red). (D) Scatterplot between Dppa2/4^+/+^ and Dppa2/4^-/-^ 2-cell embryos highlighting maternal transcripts (green) and major ZGA transcripts (purple). (E) per probe normalised heatmap showing relative expression of major ZGA transcripts in 2-cell embryos. Genotype and litter of the embryos are denoted and do not cluster together (F) Normalised expression of maternal transcripts, MERVL and major ZGA transcripts in the different embryo genotypes.

Furthermore, in the single knockouts the absence of one protein led to destabilisation of the other, consistent with what has been observed in ESCs (Eckersley-Maslin et al., 2019; Eckersley-Maslin et al., 2020). Notably, all embryos derived from maternal knockout females crossed with WT males had detectable MERVL gag protein, indicating ZGA had successfully taken place. However, both single Dppa4^+/-^ and double Dppa2/4^+/-^ embryos had lower MERVL levels than their Dppa2/4^+/+^ counterparts (Figure 3A-B) suggesting Dppa4 may have a subtle role in modulating ZGA.

To determine more globally if there are any defects in ZGA, we performed single-embryo RNA-sequencing to comprehensively survey the transcriptome in 2-cell embryos. Three matings were setup: Dppa2/4^+/+^ females with Dppa2/4^+/+^ males as controls in which all embryos would be Dppa2/4^+/+^; Dppa2/4^m-^ females with Dppa2/4^+/+^ males in which all embryos would be Dppa2/4^+/-^; and Dppa2/4^m-^ females with Dppa2/4^+/-^ males in which half the embryos would be expected to be Dppa2/4^+/-^ and half Dppa2/4^-/-^. We analysed levels of Dppa2 and Dppa4 transcripts to assign embryos from the latter category into the two embryo genotypes so they could be analysed separately (Figure S2A). Clustering analysis using t-SNE revealed that 2-cell embryos derived from Dppa2/4^m-^ females had a very similar transcriptome to those from Dppa2/4^+/+^ females (Figure 3C). Consistently, there were just 93 differentially expressed genes between Dppa2/4^+/+^ and Dppa2/4^-/-^ embryos, and only 6 genes including Dppa4 that were consistently differentially expressed across all heterozygous and homozygous embryos (Figure S2B-D). Significantly, there were no substantial differences in the expression of major ZGA transcripts between Dppa2/4^-/-^ and Dppa2/4^+/+^ embryos, indicating successful ZGA had taken place (Figure 3D-F). Moreover, maternally deposited transcripts remained unchanged (Figure 3F). In contrast to MERVL protein levels (Figure 3A-B), MERVL transcription was unaltered (Figure 3F), suggesting there may be differences in protein translation and/or stability in Dppa2/4^+/-^ embryos which warrants further investigation in future studies. Alternatively, Dppa2/4 may have more subtle effects in ZGA reflected by the immunofluorescence staining but not captured by the transcriptome analysis. Together our results here reveal that embryos from maternal knockout embryos undergo successful ZGA and were largely transcriptionally indistinguishable from those from wild type females. Therefore, maternal Dppa2/4 is not essential for ZGA *in vivo*.

### Both maternal and zygotic Dppa4 are required for offspring survival

Despite undergoing successful ZGA, maternal Dppa2/4 proteins are crucial for offspring survival. As noted above, we saw reduced litter size for Dppa2^m-^ or Dppa4^m-^ maternal knockouts which was exaggerated when heterozygous males were used (Figure 2D). Miscarriages were frequent and the offspring between maternal knockout females and heterozygous males of all three genotypes did not follow the expected 50:50 mendelian ratio for the offspring heterozygote and homozygous null genotypes (Figure 4A). Moreover, pup survival was severely impaired for both single and double knockouts with a high lethality rate by postnatal day 3 for offspring derived from maternal knockout females crossed with either wild type or heterozygous males (Figure 4B). Importantly, almost all offspring survived at similar rates to controls when wild type females were crossed with heterozygous males (in which 50% of the offspring would be expected to be heterozygous), indicating that heterozygous offspring survive if they have a functional maternal allele of *Dppa2/4*.

**Figure 4:**
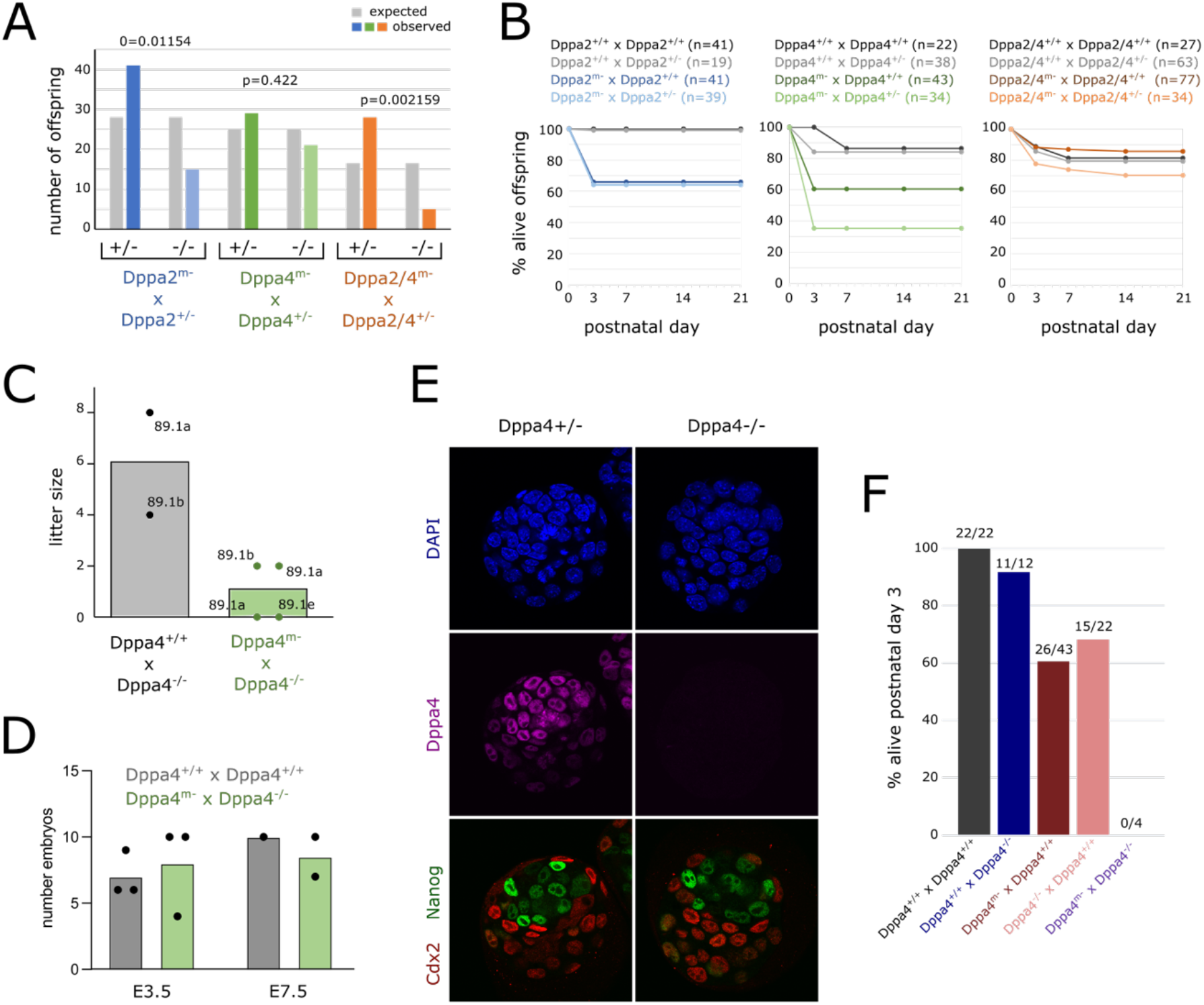
Both maternal and zygotic Dppa4 are required for offspring survival. (A) Genotype frequency of offspring (dead and alive) from Dppa2^m-^ (blue, n=56 offspring from 11 matings), Dppa4^m-^ (green, n= 50 offspring from 12 matings) or Dppa2/4^m-^ (orange, n=36 offspring from 9 matings) females crossed with the respective heterozygous males. Grey bars represent expected mendelian ratios. Dppa2 and Dppa2/4 difference is significant (Chi-Square test). (B) Percentage of live offspring at postnatal day 0, 3, 14 and 21 for wildtype females (grey) and maternal knockout females (coloured) crossed with wild type males (dark) or heterozygous males (light) for Dppa2 (blue, left), Dppa4 (green, middle) or Dppa2/4 (orange, right) alleles. Number of live pups at day 0 is denoted for each cross. (C) Litter size (dead and alive) for Dppa4^+/+^ (grey) or Dppa4^m-^ (green) females crossed with Dppa4^-/-^ males. Bars represent average. Male IDs used in matings are denoted. (D) Number of embryos collected at E3.5 or E7.5 from Dppa4^+/+^ females crossed with Dppa4^+/+^ males (grey, embryos are Dppa4^+/+^) or Dppa4^m-^ females crossed with Dppa4^-/-^ males (green, embryos are Dppa4^-/-^). (E) Immunofluorescence staining of Dppa4^+/+^ or Dppa4^-/-^ embryos collected at E3.5 for Dppa4 (magenta), Cdx2 (red), Nanog (green) counterstained with DAPI nuclear stain. Scale bar 20µm (F) proportion of offspring alive at postnatal day 3 for heterozygous Dppa4^+/-^ offspring when the null allele is inherited paternally (blue) or maternally (red, pink). No live offspring were born from crosses between Dppa4^m-^ and Dppa4^-/-^ animals. Fraction above each bar represents total live offspring at day 3 over total number of offspring born.

Despite the high lethality, we were able to obtain 3 Dppa4^-/-^ adult males and 2 Dppa4^-/-^ females from a single litter from 8 matings. No homozygous null animals survived weaning from *Dppa2* or *Dppa2/4* genotypes. We first assessed the fertility of the surviving adult Dppa4^-/-^ males by mating them to WT females where they gave rise to viable offspring (Figure 4C) indicating they were fertile. We then set up 4 separate matings with Dppa4^m-^ females which had successfully given litters in other matings. Of these, 2 resulted in miscarriages and 2 produced litters with 2 offspring each which were stillborn or died immediately after birth (Figure 4C).

We next sought to determine when during embryonic development this lethality occurred. We readily isolated similar numbers of Dppa4^-/-^ blastocysts as those from crosses between wild type females and males (Figure 4D). The Dppa4^-/-^ blastocysts did not show any morphological abnormalities with proper cavity formation, similar number of blastomeres and correctly segregated Cdx2^+^ trophectoderm and Nanog^+^ inner cell mass (Figure 4E, S3A-C). Moreover, we were able to isolate normal looking E7.5 embryos from Dppa4^m-^ x Dppa4^-/-^ crosses (Figure 4D), at a time that Dppa2/4 are no longer expressed. Together this suggests that the embryonic lethality and defects occur following implantation at a time that Dppa2/4 are no longer expressed. This is consistent with the findings from zygotic knockout mouse models (Madan et al., 2009; Nakamura et al., 2011), and a molecular function for Dppa2/4 as epigenetic priming factors where they establish a permissive epigenome to facilitate future cell differentiation (Eckersley-Maslin et al., 2020; Gretarsson and Hackett, 2020).

Lastly, we disentangled the relative importance of maternal vs zygotic Dppa4 in embryonic survival. If only embryonic levels of Dppa4 were important for embryonic development and survival of Dppa4^+/-^ animals, then it should not matter whether the mutant allele was inherited maternally or paternally. However, we only observed a marked impairment in the survival rate of Dppa4^+/-^ embryos when the mutant allele was inherited from the mother (Figure 4F). Therefore, maternal stores of Dppa4 are important for offspring survival, in addition to what is embryonically transcribed.

## Discussion

Here, we systematically and comprehensively assess the importance of maternal Dppa2/4 using single and double conditional knockout mouse models. In contrast to predictions from *in vitro* studies, maternal Dppa2/4 is dispensable for ZGA and preimplantation development *in vivo*. However, maternal Dppa4 is important for offspring survival, and the complete absence of both maternal and embryonic Dppa4 is not compatible with development.

The 2C-like cell *in vitro* system has widely been used to gain insights into the biology of 2-cell embryos and ZGA (reviewed in (Iturbide and Torres-Padilla, 2020)). However, regulators of ZGA-associated transcripts *in vitro* do not always validate as ZGA regulators *in vivo*. Recently, the transcription factor Dux was also shown to be dispensable for ZGA *in vivo* (Bosnakovski et al., 2021; Chen and Zhang, 2019; De Iaco et al., 2020; Guo et al., 2019), despite being essential for the 2C-like state *in vitro* (De Iaco et al., 2017; Hendrickson et al., 2017). Therefore, while a useful tool, it is critical that findings using the *in vitro* 2C-like cell system are validated *in vivo*.

In ESCs, Dppa2/4 function directly upstream of Dux to induce its expression (De Iaco et al., 2019; Eckersley-Maslin et al., 2019; Yan et al., 2019). Similar to our observations with Dppa2/4, Dux knockout mice also undergo successful ZGA yet have reduced litter sizes and heterozygous crosses deviate slightly from mendelian frequencies (Bosnakovski et al., 2021; Chen and Zhang, 2019; De Iaco et al., 2020; Guo et al., 2019). However, in contrast to what we observe for Dppa2/4, Dux homozygous null animals are readily obtained, indicating that Dppa2/4 has additional roles in embryo development beyond regulating Dux.

Maternal knockouts of the majority of ZGA regulators lead to a delay and/or dampening of ZGA transcription, rather than a complete absence of ZGA in its entirety (reviewed in (Eckersley-Maslin et al., 2018)). One possibility is that there is a high degree of redundancy between ZGA regulators to ensure this key developmental milestone is reached and not leaving it vulnerable to a single mutation. Alternatively, defects in regulating ZGA may not always lead to major transcriptional changes in the 2-cell embryo but may manifest more subtly with phenotypes only seen at later developmental or post-natal stages. Consequently, assessing phenotypes of maternal regulators of ZGA may be more complex than previously thought.

Importantly, our findings indicate paternal and maternal Dppa2/4 are not equal. We were able to obtain a small number of Dppa4^-/-^ males which enabled us to disentangle the importance of the maternal vs paternal alleles. Heterozygous Dppa4^+/-^ animals that inherit a maternal null allele (from maternal knockout females) have much poorer survival rates than those that inherit a paternal null allele. It remains to be determined when during development the maternal stores are important. We were able to collect morphologically normal blastocysts suggesting that the embryonic defects that lead to high mortality rate occur following implantation at a time when Dppa4 is no longer expressed, although we cannot rule out the phenotypes are not a consequence of subtly affecting ZGA. We hypothesise that maternal Dppa2/4 may be bookmarking key developmental genes in the oocyte and zygote, priming them for future activation at later developmental stages. Consistently, in ESCs Dppa2/4 function as priming factors and are required to establish bivalent chromatin, thus facilitating effective differentiation (Eckersley-Maslin et al., 2020; Gretarsson and Hackett, 2020). Consistent with a priming function in early embryos, we reveal that both *in vitro* and *in vivo*, Dppa2/4 bind mitotic chromatin. This presents a mechanism by which Dppa2/4 could mark developmental promoters through the widespread epigenetic reprogramming that takes place in preimplantation embryos, ensuring they are appropriately activated in a timely manner at later developmental stages. Embryonically expressed Dppa2/4 may reinforce this priming function and partially compensate for lack of maternal stores. This uncoupling of when Dppa2/4 are present and the phenotypic consequences of their loss is a hallmark of epigenetic priming factors (Eckersley-Maslin, 2020). In this way, maternal proteins may function beyond ZGA and implantation, to ensure the successful development of the embryo.

## Materials and methods

### Generation of conditional knockout mice

Conditional knockout mice were designed and generated by Cyagen. Homology arms were amplified by BAC clones and introduced into targeting vectors containing Neomycin selection cassette flanked by LoxP sites. For Dppa2 and Dppa4 single knockouts, exon 2 was selected as the conditional knockout region. For the double Dppa2/4 conditional knockout, exon 2 of Dppa4 through to exon 7 of Dppa2 was selected as the conditional knockout region. Targeted C57Bl/6 mouse embryonic stem cells (ESCs) were identified by genotyping PCR and confirmed by Southern Blot. Targeted ESCs were injected into C57Bl/6 albino embryos, which were re-implanted into CF-1 pseudo-pregnant females. Founder animals were identified by their coat colour and germline transmission confirmed by breeding with Flp-deleter females and subsequent genotyping of the offspring. Conditional knockout lines were maintained by intercrossing conditional homozygous (c/c) or heterozygous (c/+) animals. To generate maternal knockout mice, female c/c mice on a C57Bl/6 background were crossed with Zp3-Cre males (Lewandoski et al., 1997). All experimental procedures were performed in accordance with the Animals (Scientific Procedures) Act 1986 and by local authority granted by the Animal Welfare and Ethical Review Body (AWERB) committee of the Babraham Institute.

### Embryo collection

All mouse embryos were collected from natural matings between the appropriate genotypes according to standard procedures (Hogan, 1994), at different timepoints, depending on the desired stage. MII oocytes were collected from C57Bl/6 females in oestrus, and zygotes, 2 cell, 4 cell, 8 cell, morula and blastocysts at E0.5, E1.5, E2, E2.5, E3 and E3.5 respectively. Embryos used in this study were derived from a cross of C57Bl/6 females mated with C57Bl/6 males. The KO embryos were derived from the maternal knockout females mated with either C57Bl/6 proven studs or the appropriate genotype (Dppa2/Dppa4/Dppa2-4^+/-^ or Dppa4^-/-^) males.

### Mouse embryonic stem cell culture

E14 mouse embryonic stem cells were cultured in serum/LIF conditions (DMEM, 4,5000mg/L glucose, 4mM L-glutamine, 110mg/L sodium pyruvate, 15% fetal bovine serum, 1U/ml penicillin, 1mg/ml streptomycin, 0.1mM nonessential amino acids, 40mM b-mercaptoethanol, 10^3^ U/ml LIF). Plasmids containing full length Dppa2-GFP and Dppa4-GFP were generated previously (Eckersley-Maslin et al., 2019). Truncations were generated by amplifying appropriate sections of the constructs and cloning into the pDONR221 vector. Gateway cloning was then used to transfer the truncated cDNA sequences into an in-house built pDEST vector as previously described (Eckersley-Maslin et al., 2019). Plasmids were transfected into E14 ESCs using Lipofectamine 2000.

### Immunofluorescence staining

Oocytes and embryos were washed in PBS, fixed for 15 min in 4% paraformaldehyde in PBS, permeabilized with 0.5% Triton X-100 in PBS for 1 h, and blocked in 1% BSA in PBS for 1 h. Primary antibodies were diluted in 1% BSA and the embryos incubated for 1 h. After 1h wash in 1% BSA, the samples were incubated in secondary antibodies for 45 min, followed by a 1 h wash in PBT (0.05% Tween20 in PBS). DNA was counterstained with 5 ug/ml DAAPI in PBS, and embryos mounted in fibrin clots. All incubations were performed at room temperature. Primary antibodies and dilutions used were: mouse anti-Dppa2 (Millipore, mab4356) 1:100, goat anti-Dppa4 (R&D Systems, AF3730) 1:100, rabbit anti-MERVL (Huabio, R1501-2) 1:200, mouse anti-Cdx2 (Biogenex, MU392-UC) 1:200, rabbit anti-Nanog (Abcam, ab80892) 1:200. Secondary antibodies used were anti-rabbit AF-conjugated 647, anti-mouse AF-conjugated 568 and anti-goat AF-conjugated 488 (Molecular Probes) and diluted 1:1000. Single optical sections and Z-stacks were captured with a Zeiss LSM780 microscope (63x oil-immersion objective). Fluorescence colocalization analysis was performed with ImageJ, and fluorescence intensity measurements were performed with Volocity 6.3 (Improvision). The nuclei of each blastomere in the 2 cell embryos were measured separately and their average was used for the final analysis. The plots were generated using RStudio.

### Single-cell RNA-sequencing

Embryos from the Dppa2/4^m-^ were generated from natural matings between males and females of relevant genotypes. The zona pellucida was removed using Tyrode’s solution (Sigma, T1788), and individual embryos were placed in in 2.5µl methyltransferase reaction mixture, according to the published protocol (Clark et al., 2018). mRNA was captured using Smart-seq2 oligo-dT pre-annealed to magnetic beads (MyOne C1, Invitrogen). The lysate containing the gDNA was transferred to a separate PCR plate, and the beads were washed 3 times in 15 ml FSS buffer (Superscript II, Invitrogen), 10 mM DTT, 0.005% Tween-20 (Sigma) and 0.5 U/ml of RNAsin (Promega). The beads were then resuspended in 10 ml of reverse transcriptase mastermix (100 U SuperScript II (Invitrogen), 10 U RNAsin (Promega), 1 X Superscript II First-Strand Buffer, 2.5 mM DTT (Invitrogen), 1M Betaine (Sigma), 9mM MgCl2 (Invitrogen), 1 mM Template-Switching Oligo (Exiqon), 1 mM dNTP mix (Roche)) and incubated on a thermocycler for 60 min at 42°C, followed by 30 min at 50°C and 10 min at 60°C. PCR was then performed by adding 11 ml of 2 X KAPA HiFi HotStart ReadyMix and 1 ml of 2 mM ISPCR primer, and cycling as follows: 98°C for 3 min, 15 cycles of 98°C for 15 sec, 67°C for 20 sec, 72°C for 6 min, and finally 72°C for 5 min. cDNA was purified using a 1:1 volumetric ratio of Ampure Beads (Beckman Coulter) and eluted in 20 ml of water. Libraries were prepared from 100 to 400 pg of cDNA using Nextera XT Kit (Illumina), per manufacturer’s instructions but with one-fifth volumes for each sample. Libraries were sequenced on an Illumina NextSeq500 MidOutput 75 bp paired-end reads per embryo.

### RNA-sequencing analysis

Libraries were trimmed using Trim Galore (v0.6.5, Cutadapt v2.3) and mapped to the mouse GRCm38 genome assembly using HISAT2 (v2.1.0, --no-softclip) and filtered to have MAPQ scores of 20 and above. Data was quantified using the RNA-sequencing quantitation pipeline in SeqMonk (https://www.bioinformatics.babraham.ac.uk/projects/seqmonk/), without strand-specific quantification, using mRNA probes. All the analysis, specified in each figure legend, was carried in Seqmonk, and only the violin plots were generated in RStudio. The major ZGA and maternal deposited gene lists used in the analysis were generated using the publicly available dataset (GSE44183) from (Xue et al., 2013).

## Acknowledgements

We would like to thank all members of the Reik laboratory for their helpful discussions and Federico di Tulio for his assistance with cloning. We also thank Jasmin Stowers for her mentorship of OK, and Wendy Dean and Celia Alda-Catalinas for helpful advice on mouse genetics, breeding and colony management. We thank all staff in the Babraham Biological Support Unit (BSU), Christel Krueger, Felix Krueger and Simon Andrews in the Bioinformatics facility, Simon Walker and Hanneke Okkenhaug in the Imaging Facility and Nicole Forrester and Paula Kokko-Gonzales in the Sequencing facility at Babraham Institute for their support. Research in the Reik lab is supported by the Biotechnology and Biological Science Research Council (BBSRC BBS/E/B/000C0421) and the Wellcome Trust (210754/Z/18/Z). O.K. is supported by a MRC-DTP PhD Studentship and M.A.E-M was supported by a BBSRC Discovery Fellowship (BB/T009713/1).

## Competing interests

The authors declare that they have no competing interests.

## Author contributions

MAE-M, WR and OK conceived and designed the study. OK and MAE-M maintained the mouse colonies. OK, FS and DD carried out the embryo collections. OK performed immunofluorescence staining, imaging and IF data analysis. OK generated single-cell RNA-sequencing libraries and OK and MAE-M performed bioinformatic analysis. MAE-M and OK wrote the manuscript. MAE-M and WR supervised the study.

## Notes

### Competing Interest Statement

The authors have declared no competing interest.

